# Habituation, task solving and memorization may facilitate biological invasions: the Starling example

**DOI:** 10.1101/2021.07.24.453664

**Authors:** Alexandra Rodriguez, Martine Hausberger, Philippe Clergeau, Laurence Henry

## Abstract

Invasions ecology deals more and more with behavioural characteristics of invasive species. Particularly, research have focused on the personality of invaders and on their way of coping with novelty in new habitats. Traits of neophobia may limit individuals in their exploration of novel objects or the consumption of novel foods, they may stop the access to valuable ressources. Actually, in novel environments like cities, food can be unreachable in throwaway dishes with lids or hidden in the garbage or even close to frightening objects. Animals may either left the place and waste these resources, or they can express low neophobia from the beginning and manipulate the objects to reach food. They may also habituate progressively to the context and use the ressources.

Here we analyzed the behavioural responses of individuals from three populations of European starling *Sturnus vulgaris*: a population anciently settled in a rural region, a population that has recently colonized a urban area and a population of wintering migrant birds.

We used a series of tests in order to explore if individuals would habituate to a novel object and if they could remember it eight months later. We explored if individuals would be less neophobic when confronted to two novel objects successively and we tested them in a learning task involving a novel object and an attractive food. Our results show that *Sturnus vulgaris* habituates rapidly to novel objects and that categorization facilitates neophobia lost when confronted to two different novel objects. Initially young birds appeared to be more skilled than adults in the learning task. Individuals from this species seem to be able to remember an object durably. We suggest that habituation, task solving and memorization are three mechanisms enhancing biological invasions.

## Introduction

When they arrive into a novel environment, animals may be confronted with several novel stimuli as novel food, novel predators or even novel sounds as each landscape has its particular comunities and its particular soundscape (Rodriguez et al. 2014). Animals scan also be confronted with new objects and new problems to reach food. Approaching novel objects and having access to the food may imply an attenuation of neophobia towards the objects and expressing particular behaviours in order to overcome the obstacles to reach the food. Behaviours as approaching, exploring and manipulating objects are often involved in these processes.

If animals arrive into habitats containing many different objects it may be necessary to be able to tolerate these objects durably and to habituate to their permanent presence. This can be particularly the case of habitats like urban areas where several coloured and mobile objects are present (Seferta et al. 2001, Echeverria et al. 2006).

Hesitancy to approach novel objects and thus to frequent and manipulate them varies across the species (Mettke-Hofmann et al. 2002) and across the ecological contexts in which animals live. It has been demonstrated that levels of neophobia towards new objects and there subsequent implications in learning vary even within a same family as in the taxa of parrots, sparrows and warblers (Greenberg 2003).

European starling, *Sturnus vulgaris* has reproduced and expanded with success in several of the regions it has been introduced as North America, Australia, South Africa, New Zeland and Argentina (Pell and Tidemann 1997, Peris et al. 2005).

In a previous study we analyzed interindividual differences of neophobia towards a novel object in starlings from three different populations: a migratory population of the northeastern regions of Europe, a population of a rural area in Brittany where starlings have been established for at least 400 years and a urban population from an area in Brittany colonized 80 years ago by the species. In this study we found that there were more neophobic individuals than bold individuals in the rural group when confronted to a novel object (Rodriguez et al. 2018).

However, overcoming fear and approaching food that can be close to novel objects or manipulating these objects in order to obtain food may be crucial for individuals independently of their original populations. By consequence, coping with initial neophobia may be essential to avoid starvation, particularly when they visit novel habitats or in winter when food is scarce and difficult to reach. It has been demonstrated that tits and parrots species manipulate objects like milk bottles and garbage containers indicating that they overcome their neophobia and develop cognitive strategies in order to obtain what they want (Fisher and Hinde 1949, Gadjon et al. 2006).

Other studies have demonstrated that bird species like Cattle Egrets, *Bubulcus ibis,* will feed close to active bulldozers whereas Turkey vultures, *Cathartes aura,* will not eat in the same places until the bulldozers have not left the area (Burger and Goshfeld 1983). In mammals like coyotes, *Canis latrans,* areas containing novel objects like cones artificially put for experiment would initially provoke aversion and posterior removal provokes exploratory behaviours in the same places (Heffernan et al. 2005).

Here we explored the reaction of the three previously studied groups of starlings towards different situations involving habituation, learning and memorization tasks in order to see if the birds’ behaviour would be plastic. We wondered if the birds would modulate their behaviour towards novel objects after a first experience and if it could have a long term effect on the behaviour of the individuals towards the same objects. We also wondered if these kinds of mechanisms would be expressed in similar ways between individuals from populations with different histories or if they would be qualitatively different and particularly relevant in birds from populations usually confronted with novel environments.

## Methods

### Subjects and housing

#### Adult migratory groups

21 adult starlings (eleven males and ten females) were captured during autumn 2006 with nets at Normand cliff of Etretat. These individuals belonged to northeastern European populations that hibernate in France during winter. The individuals were housed in an aviary during one year (until autumn 2007).

Ten naïve adults (five males and five females) (group 1) were used to assess the changes in behaviour when exposed to two successive novel objects. The other eleven adults (six males and five females) (group 2) were used to analyse habituation towards a novel object and in the learning test.

#### Young groups

We removed 38, 5-14 day old nestling starling from 32 nests of two natural populations (20 chicks from rural Breton population and 18 chicks from urban Rennes population). Two series of captures were conducted: one in spring 2007 and another in spring 2008.

The 22 individuals captured in May 2007 and the 16 individuals captured in May 2008 were reared and housed in the same way. They were hand-raised using commercial pellets (Végam, Grosset) mixed with water. Before fledging, they were put in plastic boxes with wood separations (5-6 chicks per compartment) and then they were put in small cages when fledging.

Urban and rural individuals were reared together. They were hand-raised during five weeks until they could eat alone. At the age of two months they were put in an out-door aviary where they had natural sunlight and where they could hear adult starlings singing in the area. At the age of six months they were transferred to an in-door aviary were they were housed until the start of the experiments.

All the young individuals were tested for the learning task at the end of October (2007 and 2008) when they were 7 months old. The young individuals tested in 2007 were used for memorisation experiments eight months later in July 2008 when they were 15 months old.

### Behavioural experiments

For the tests we used 120 x 40 x 50 cm cages that can be divided in two identical parts by a plastic opaque separation.

Exposure to two successive novel objects:

The novel object tests consisted in putting an unknown object in the second part of the cage and to observe the reaction of the individual. At the beginning of the test the bird was in the first part of the cage. The observer put the new object in the second part and removed the plastic separation between the two parts of the cage.

There were four perches in the cage. There were a feeding dish and drinking trough in the first part of the cage. The test lasted 15 minutes.

The first day the group of ten migratory adults was exposed to a first object (Object 1) which consisted in a Petri box with brown scotch around it and fixed to a wooden white support (Figure 1). One day after this first experiment, the individuals were confronted in the same way with a second object (Object 2) which was also unknown for them. This object was taller than the first one and consisted in a white opaque box fixed on a white wooden support (Figure 1).

**Figure 1:**
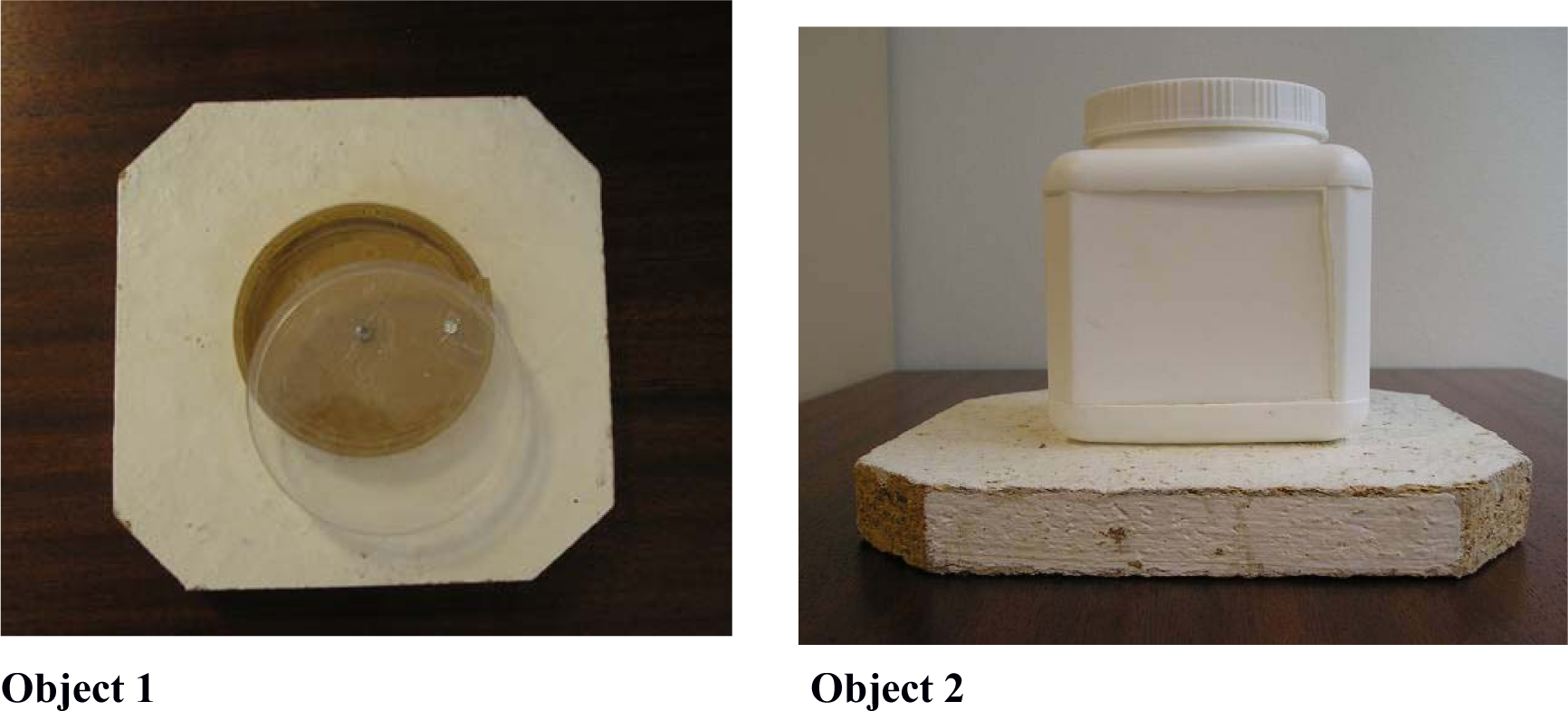
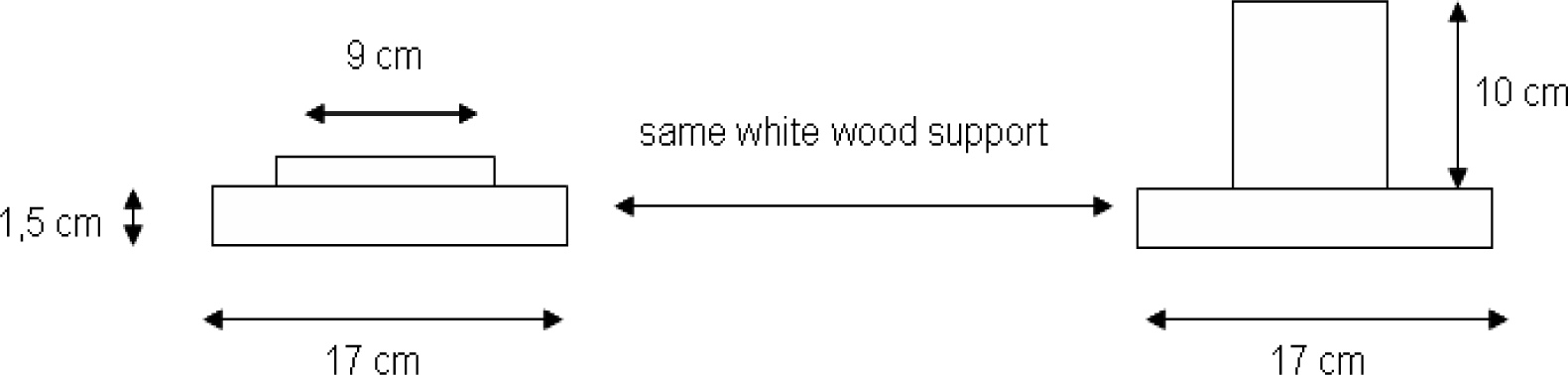
Objects used in the successive novel object tests

During the two novel objects experiments we noted all the behaviours performed by the individuals using focal sampling observations. The different behaviours expressed by the birds are listed in Table 1 and the chronology of the experiments is presented in Figure 2.

**Figure 2:**
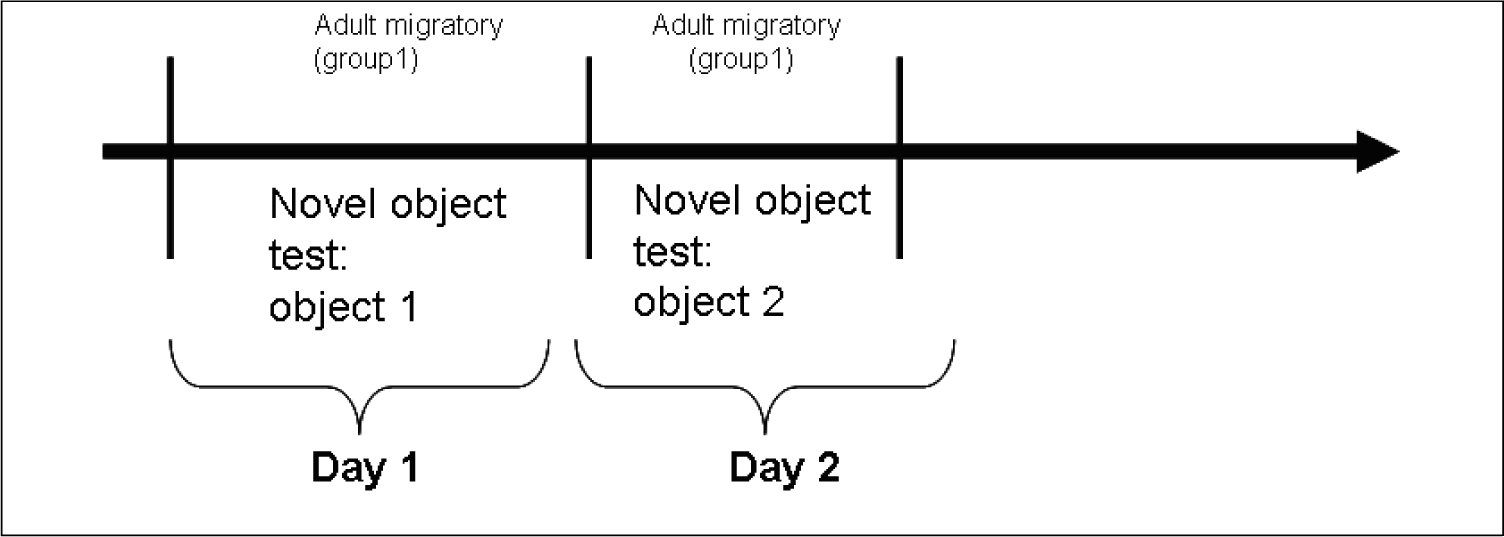
Chronology of the two novel objects successive experiments

Habituation analysis:

In a previous study, we had analysed interindividual differences in neophobia towards the novel object 1 in the group 2 of adults and on the groups of rural and urban young. During this test we had measured the latency to approach the object (latency to enter the ground of the second part of the cage). After the novel object test the Object 1 was let in the cage during the 24 hours following the novel test. For the habituation analysis we compared the latency to approach the object during the first encounter with the object and the latency to approach it during the learning task conducted one day after (Figure 3).

**Figure 3:**
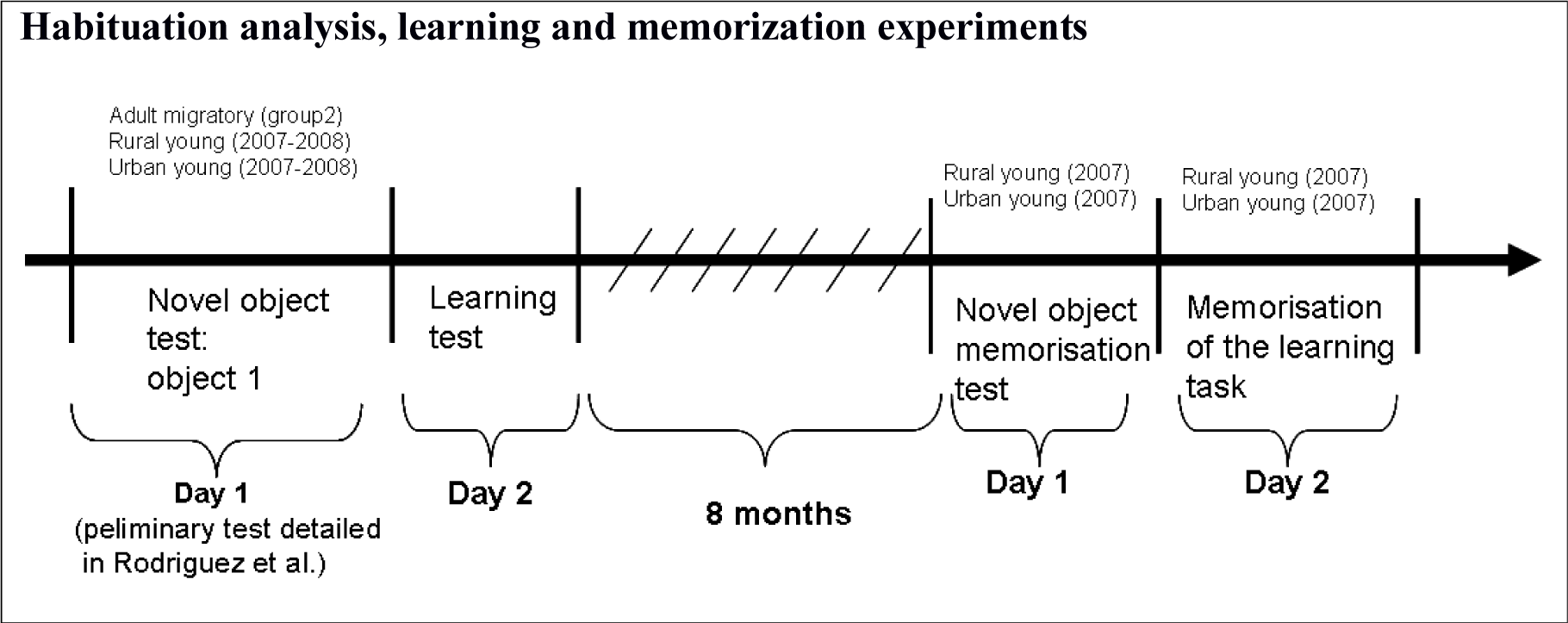
Resume of the experiments conducted on each group

### Habituation analysis, learning and memorization experiments

Learning experiments:

This test consisted in putting a worm of flour (very appreciated by Starlings) in the Petri box and covering it with its plastic lid. The plastic lid was put back to front and fixed by a nail to the wood support in order to allow it moving horizontally and get open. The experimenter put the worm in the Petri box in a way the bird could see the worm and closed the box. Then the experimenter removed her hand from the cage and started the chronometer at the moment she left the cage. She noted the latency between the moment she left and the moment the bird opened the box. The experiment lasted 15 minutes if the bird did not open the box and it was stopped at the moment the bird opened it.

The experimenter also noted the latency to touch the object for the first time (TC1, Figure 4) in order to see if there was still neophobia towards the object, and the total time (TT1) the bird manipulated the object with his beak or touched with its leg during the experiment. The observer also noted the number of blows of beak the bird made as attempts to open the box.

**Figure 4:**
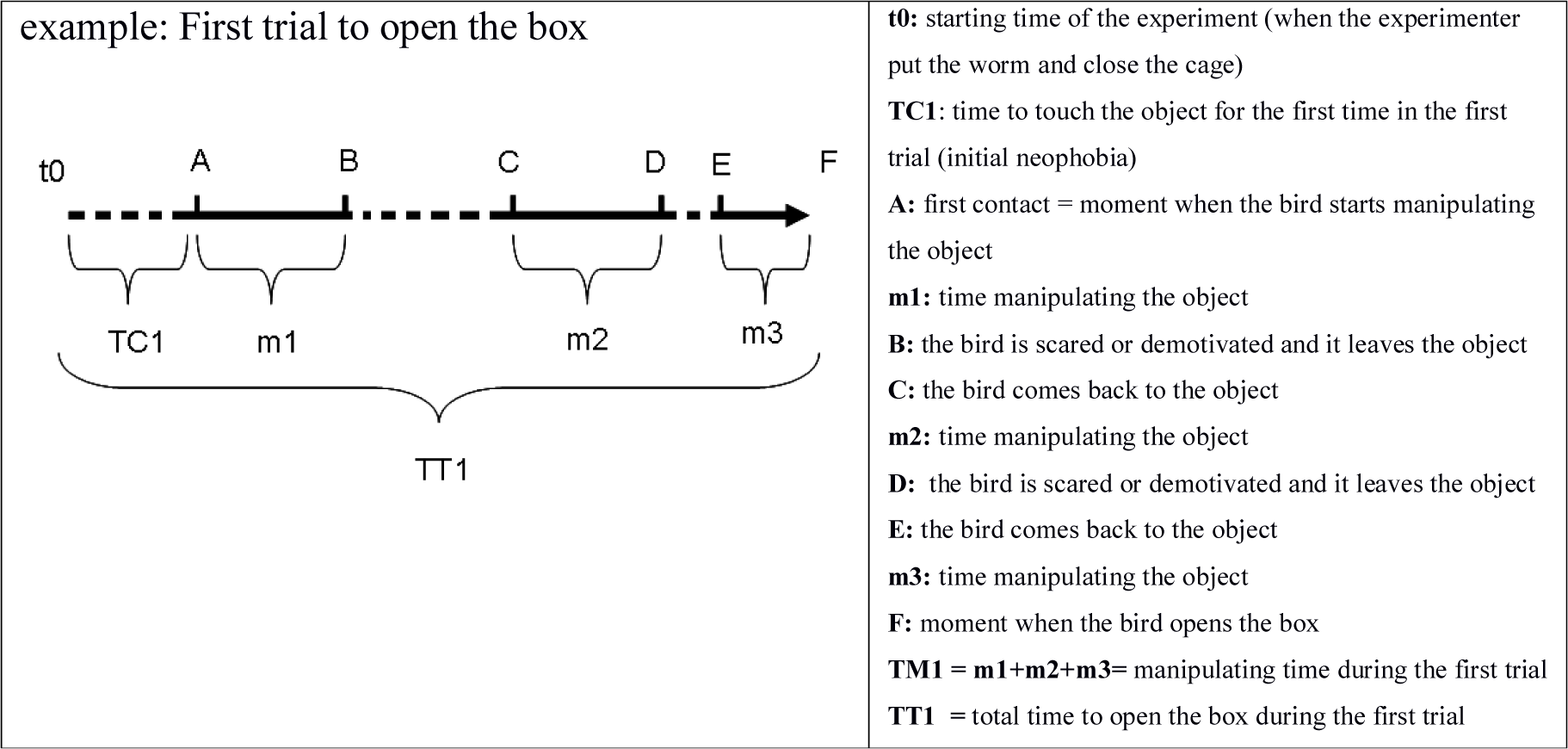
Time parameters measured during one trial of the learning task

Three experiments of this kind were conducted in order to evaluate the learning process of birds and to see if the individuals improved in the task.

For each trial, we measured the latency to touch the object for the first time, the total latency to open the box, and the time spent manipulating the object as it is presented in Figure 4.

Memorization of the object:

18 young birds tested in 2007 for the novel object test were tested again in 2008 in an identical experiment in order to see if they remembered the object.

Memorization of the solving task:

The same 18 young birds were also tested again on the task consisting in opening the box in order to see if they behave as individuals that never learned how to do the task or to see if they did better than the very first time.

### Statistical analysis

We conducted a regression analysis in order to see if the latency to touch the object in the learning task was correlated with the latency to approach the object or to the latency to touch it in the novel object test.

We conducted Cox survival models implemented in R.2.8.1 in order to test if there was an effect of sex, of the origin or of the year of capture on the young birds’ probability to open the object during the experiment. We also conducted this kind of test into the migratory group in order to see if there was an effect of the sex in adults. Finally, we conducted Cox analysis to test if there was a difference in the probability to open the box during the test between all the young and the adults.

For individuals who could open the box we tested for potential improvement of the task in the two following trials.

As some individuals improved the opening task at the second trial and others at the third trial when comparing to the first one, we conducted paired Wilcoxon tests between the first trial latencies and the following trials as between the first trial and the minimum latencies of second or third trial (the better performance between the two following trials was kept).

For testing if there was a memorisation of the object, we conducted paired Wilcoxon tests between the number of times each behaviour was expressed during the first encounter with the object and the number of times each behaviour was expressed during the 15 minutes of test conducted eight months later.

For testing if there was a memorisation of the opening task, we compared the latencies to touch, manipulate and open the box between the first trial conducted in 2007 and the trial conducted in 2008. We conducted Wilcoxon tests comparing these latencies. We also tested the number of blows of beak given in each trial.

## Results

### Modifications of reaction in successive confrontation with two novel objects

When the birds were confronted to the Object 2, the occurrences of several behaviours were maintained (Wilcoxon p>0.05). Interestingly, the frequency of gazes at the object was similar for Object 1 and 2 indicating that visual attention was equivalent in both cases probably because the two objects were new and different and they needed to be identified (Figure 5). On the contrary, the frequency of mobility behaviours decreased (flying and walking occurrences) but only the frequency of fights was significantly lower in the experiment with Object 2 (p=0.047). General observation and maintenance behaviours were equivalent from one test to the other.

**Figure 5:**
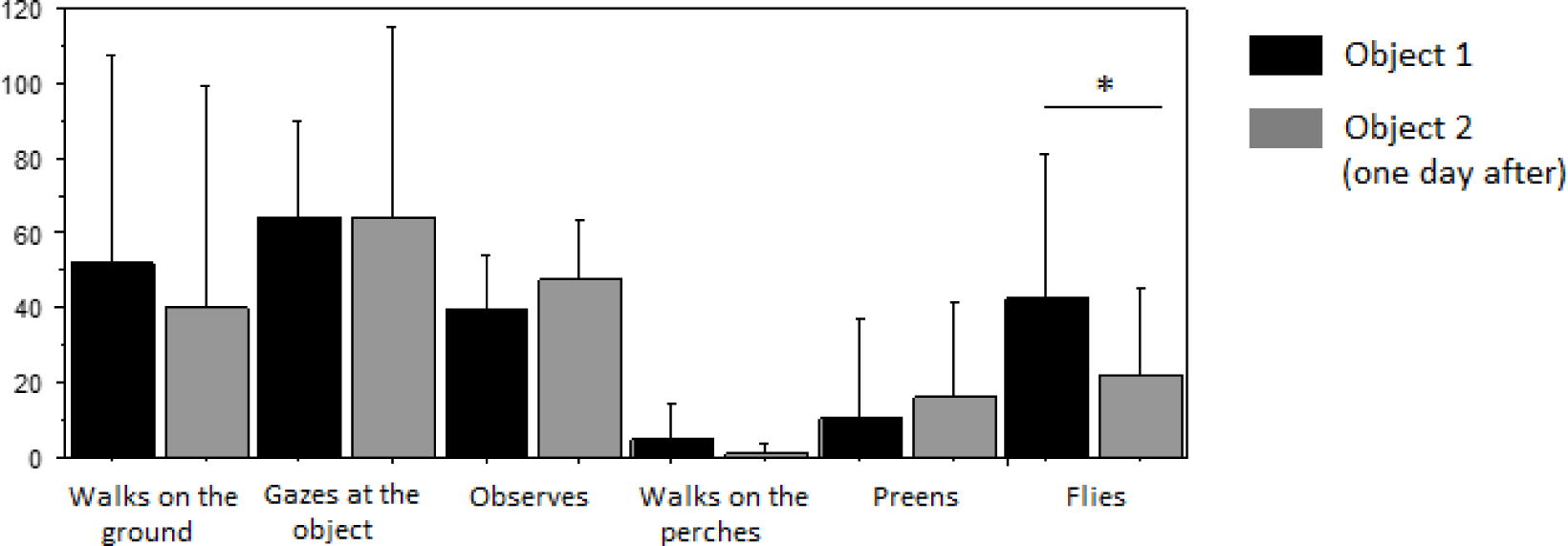
Behavioural frequencies when starlings were confronted to the two novel objects

### Comparison between initial object neophobia and first contact with object in the learning task

When we compared the latency to approach the novel object in the neophobia test and the latency to touch the object for the first time in the learning task we obtained that there was no correlation between these two latencies (R=0.22 df = 46 p>0.05). Concerning the regression between time to first contact in neophobia test and time to first contact in the learning task we obtained (R= 0.17 df= 46 p>0.05). In fact almost all the individuals seemed to have been habituated to the object during the 24 hours of its presence (Figure 6) or to be sufficiently motivated by the flour worm to overcome their previous neophobia.

**Figure 6:**
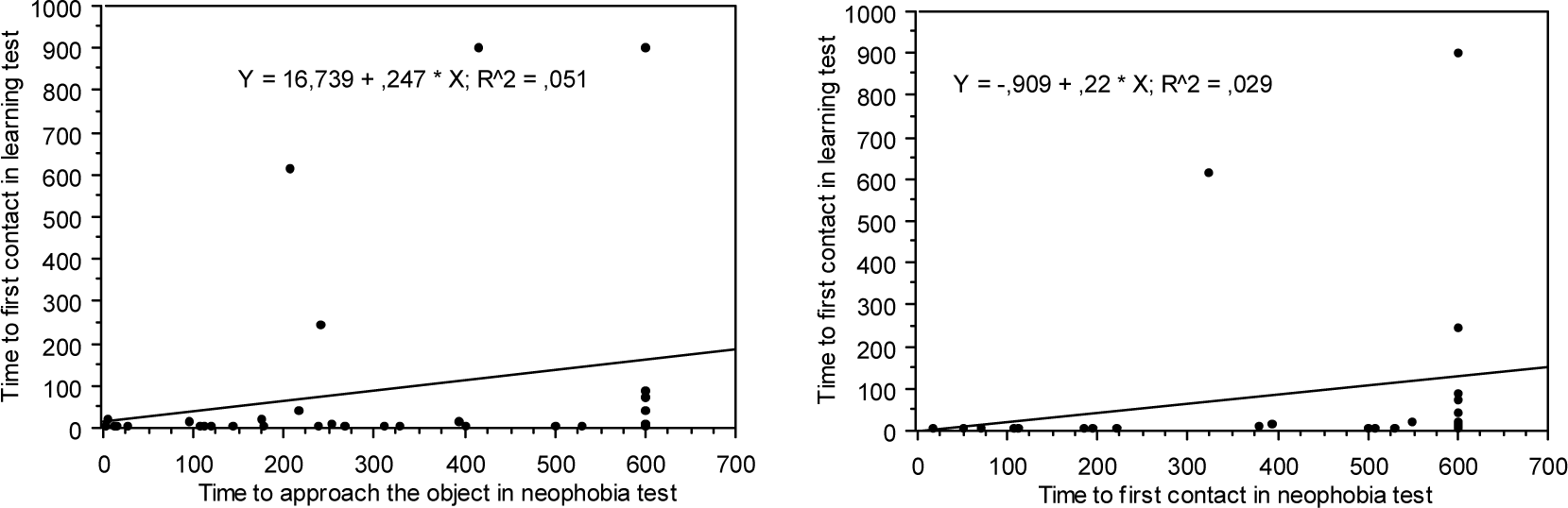
Regression and correlation values of latencies to approach and contact the object between neophobia test and learning experiment

The latency to touch the object in the learning task was inferior to one minute for mostly of them (even for some individuals that did not approach the object during the neophobia test).

Binomial tests conducted on the number of individuals who approached the object and those who did not approach it during the neophobia test resulted in a p value of 0.19 indicating that the proportion of individuals approaching the object during the neophobia test was not significantly different from 50% (Figure 7). In the same way, for the number of birds touching the object we obtained a p value of 0.11 indicating that the proportion of birds touching the object in the neophobia test did not differ from 50%.

**Figure 7:**
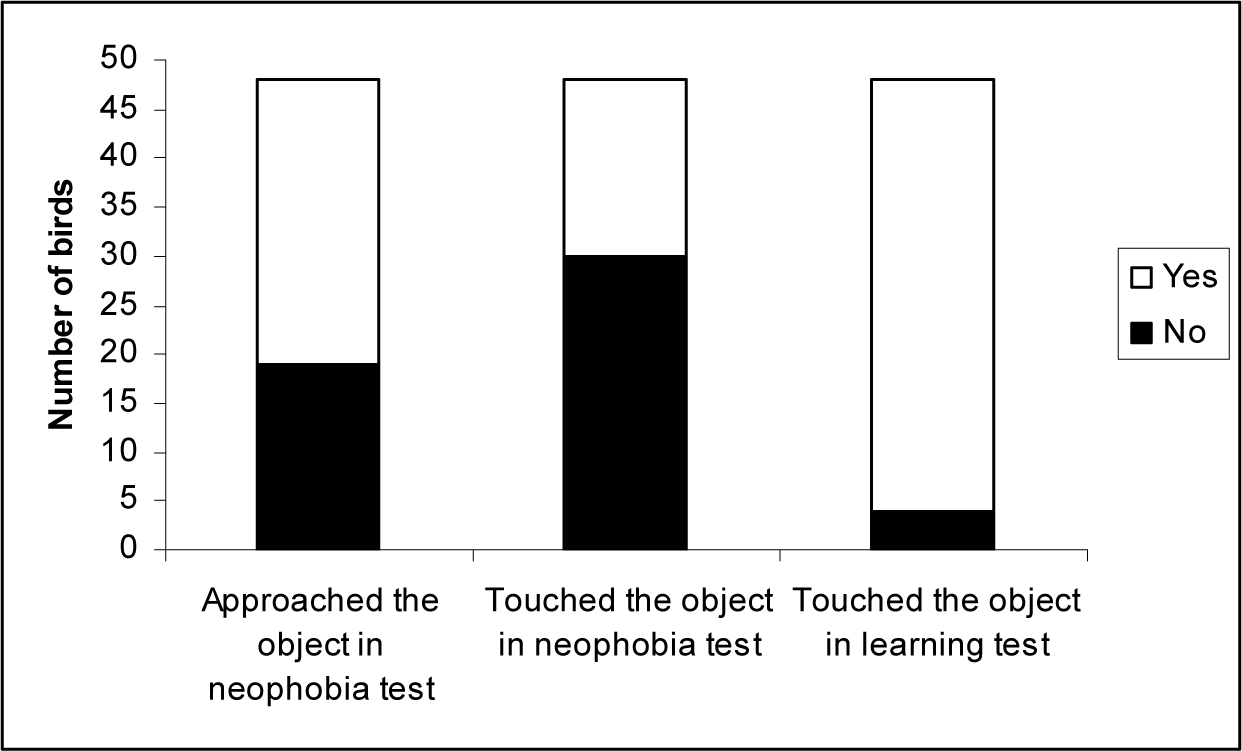
Number of birds who approached and touched the object in the tests

On the contrary, for the learning task we obtained that the proportion of birds touching the object was significantly higher than 50% (p<0.0001). This result converges with the information given by the comparison of the latencies to touch the object.

### Learning task

As we mentioned before, almost all the individuals were able to open the box. Only six individuals of the 48 tested did not open the box.

The individuals that could not open the box were: two urban young, two rural young and two migratory adults. One urban young never touched the object. The other urban young touched 17 times the object with its leg but he expressed behaviours of neophobia as he used to fly away rapidly each time he approached the object. The two young rural birds that did not open the box were very neophobic and they never approached the object. Finally, concerning migratory adults, one of them never approached the object and the other one tried to open the box. He gave 128 blows of beak to the lid but he was very shy and his weak attempts to open the box revealed a moderate level of neophobia. After his various attempts, he stopped trying and seemed unmotivated.

Despite these few cases, the big majority of individuals could open the box, indicating that in most of the cases neophobia reached very low levels; motivation and attempts were enough to allow them to open the box.

### Evolution of time to touch the object, to manipulate it and to open the box

#### The whole individuals

Wilcoxon tests results indicate that individuals improved their latencies to open the box in one of the two following trials (p<0.05 for the comparison between the first trial and the minimum of the second and the third trial) (Figure 8). The improvement of the total time to open the box was significant between the first and the second trial. This improvement was due to a significant reduction of the time to contact the object between the first and the second trial indicating that there was either a significant reduction of neophobia or a significant increasing of motivation operated by the reinforcement when birds catch the worm (or a combination of both factors).

**Figure 8:**
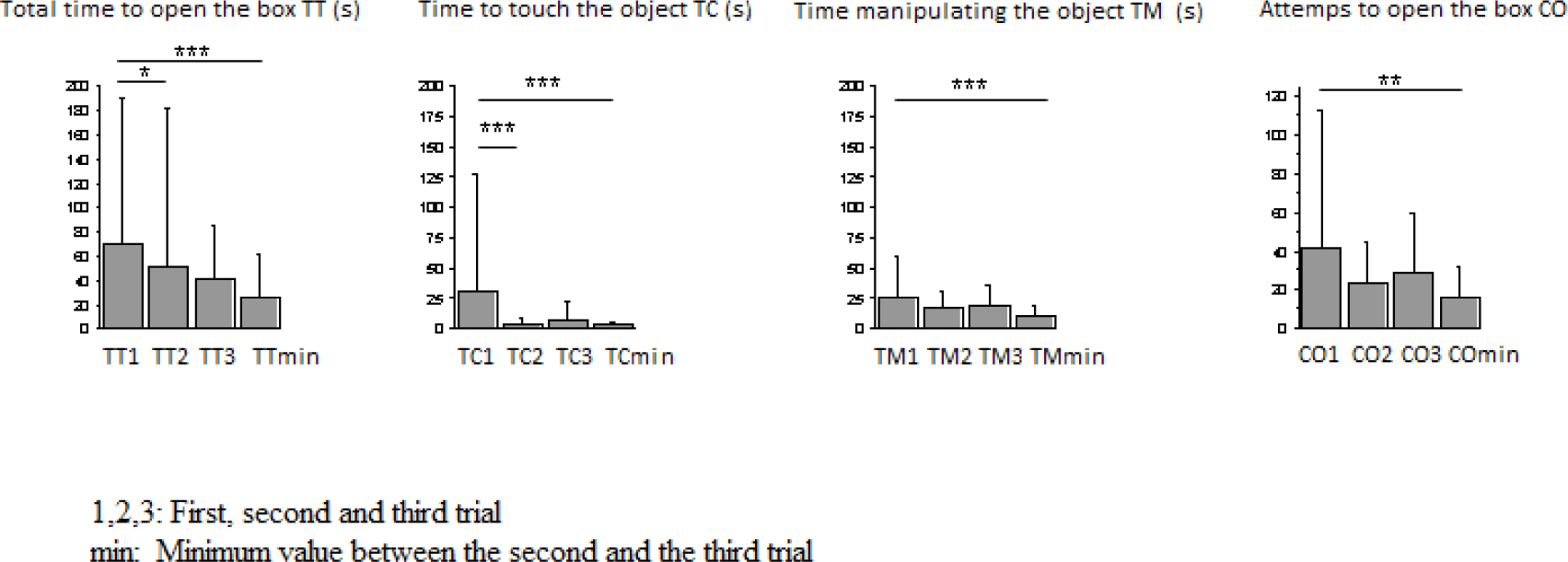
Evolution of the total time to open the box (TT), of the time to touch the object (TC), of the time manipulating the object (TM) and of the number of attempts to open the box (TO) by trial

There was also a significant decrease in manipulation time between the first trial and the best of the two following trials. The number of attempts traduced by the number of pecks on the box also significantly decreased. That means that for each individual there was at least one of the two following trials in which they manipulated better the object and the number of unfructuous attempts decreased.

#### Comparisons within and between the groups

The Cox analysis indicated that there was no effect of year of capture or sex on the probability of young starlings to open the box in any of the three trials. There was either no effect of sex on adult migratory (p>0.05 for all the Cox comparisons).

When we analyzed the total time to open the box group by group we observed that this latency was differently modified depending on the group. Total time to open the box decreased between the first and the second trial for only for the rural young group (Figure 9). The decrease of total time to open the box in rural young was due to a significant decrease in the time to touch the object and to a decrease of the number of attempts, which means that rural young both lost a part of their neophobia and improved in the opening task with experience. They also manipulate the object during a period significantly shorter in the second trial in comparison to the first trial. The urban young group tended to touch the object sooner in the second trial and improved its neophobia levels in one of the two last trials (see minimum value on Figure 9).

**Figure 9:**
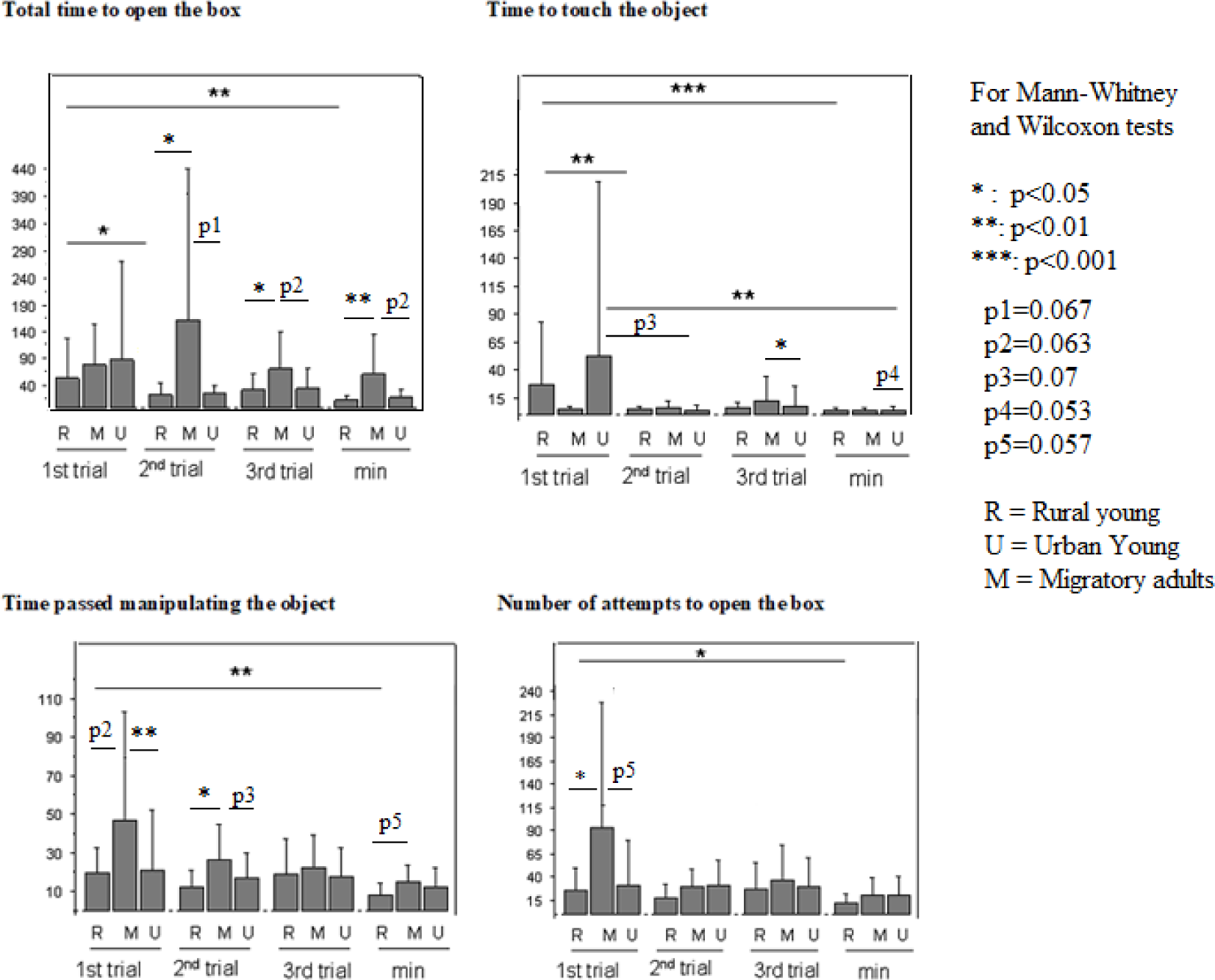
Evolution of latencies to touch, manipulate and open the box and of the number of attempts to open the box across the trials by group

Concerning the group of migratory adults, they manipulated the object longer and had to do more attemps to open the box than the young groups in the first trial. In th second trial the total time to open the box was longer in the migratory group and this was due to a longer time manipuating the object. In the third trial the tree groups took equivalent times manipulating the object but adult migratory was longer than the urban group to touch the object.

Correlation tests conducted on the latencies revealed that the total time to open the box was significantly correlated to the time to touch the object in both groups of young (Figure 10). For the rural young the total time to open the box was also significantly correlated to time to manipulate the object.

**Figure 10:**
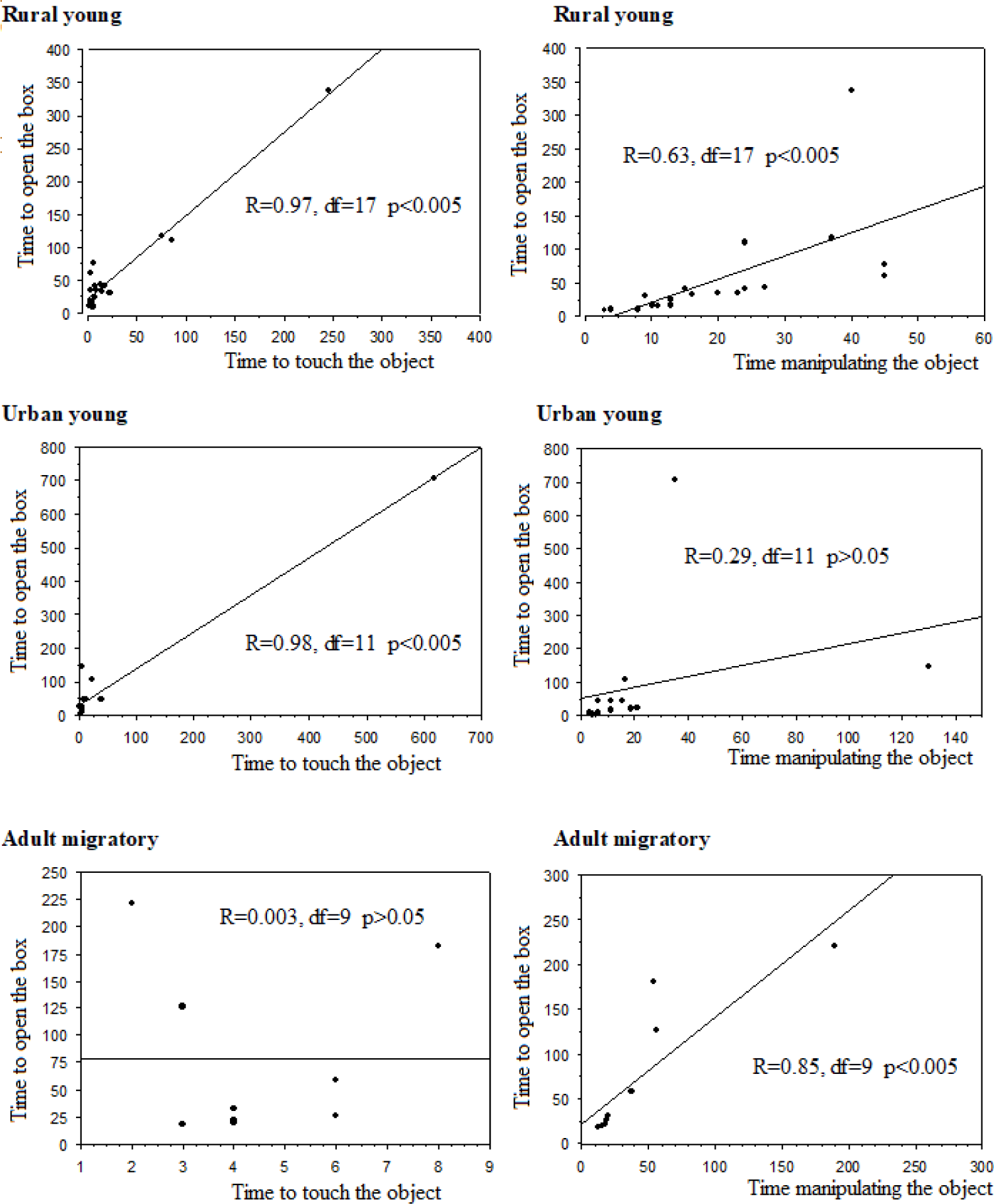
Linear regressions conducted between the total time to open the box in the first trial and time to touch and manipulate it

Finally, for migratory adults, the total time to open the box was not correlated to the time to touch the object but it was only correlated to the manipulation time. The part of variation observed in total time to open the box was explained by periods after the first contact with the object in which they did not manipulate the object. In fact there were individuals who left the object several times either because they were demotivated, or tired or even because they still expressed hesitancy. They also flew away when they were scared by abrupt noisy movements they provoked when manipulating the lid.

### Memorisation of the object

The frequency of several behaviours was modified between the first time the individuals were confronted to the Object 1 and the second time they saw the object eight months later. These results indicate that the second time they did not behaved as if they had never seen the object.

Individuals emitted significantly more calls in the second test (Figure 11, Wilcoxon test p<0.05).

**Figure 11:**
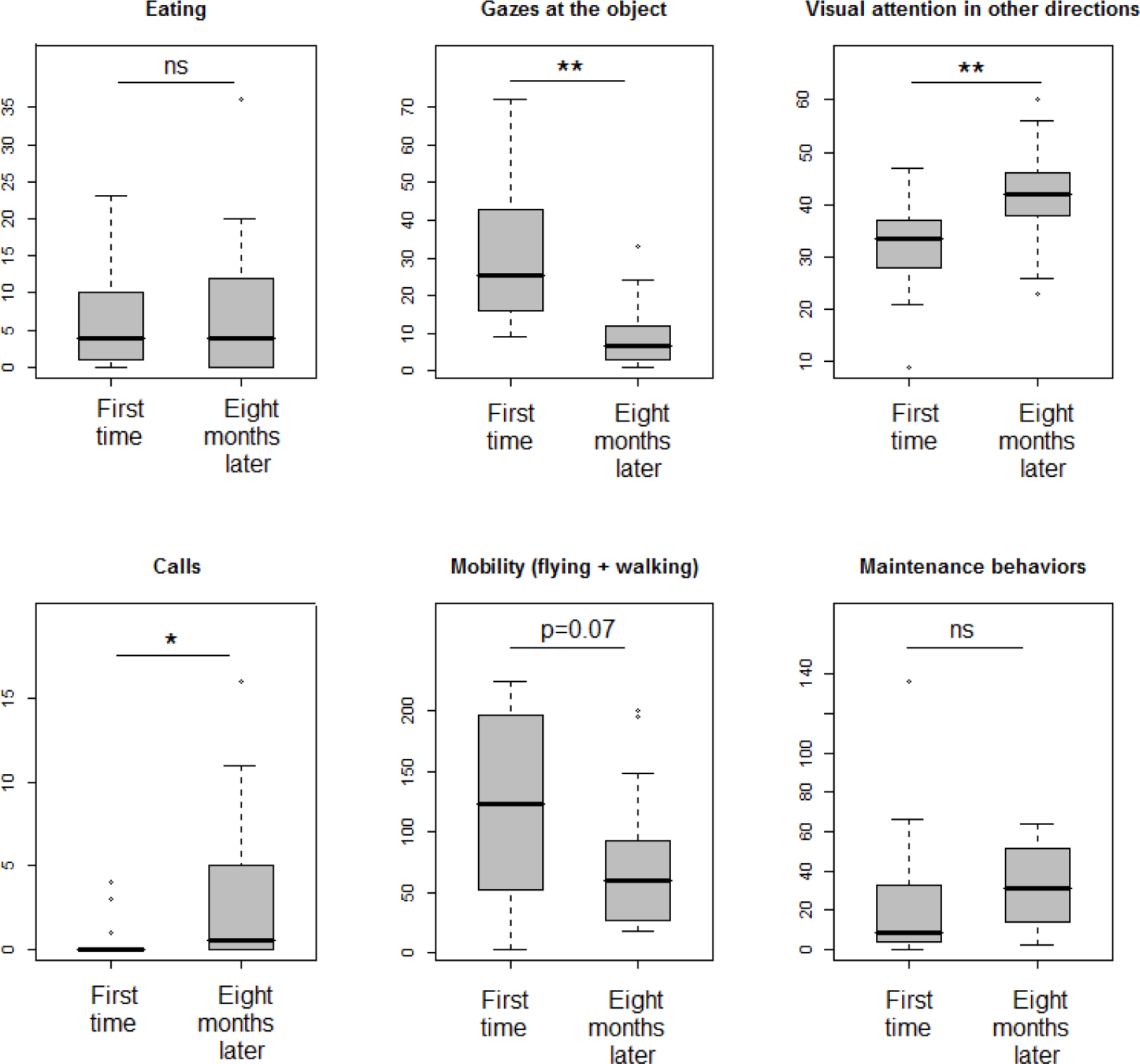
Frequencies of the behaviours during the first encounter with the object and during the second encounter eight months later

Visual attention was modified. During the first experiment, individuals gazed at the object significantly more times than during the second encounter (Wilcoxon test p<0.01). On the contrary, their visual attention was more frequently directed to other directions in the second test than in the first one (Wilcoxon test p<0.01).

The mobility tended to decrease, between the first and the second test (Figure 11).

### Memorization of the task

When we compared the first trial to open the box and the trial eight months later we obtained that manipulation time and the number of attempts to open the box (number of pecks on the lid) tended to decrease (Figure 12). It seems to be a reluctant memory of the way of solving the task after this period.

**Figure 12:**
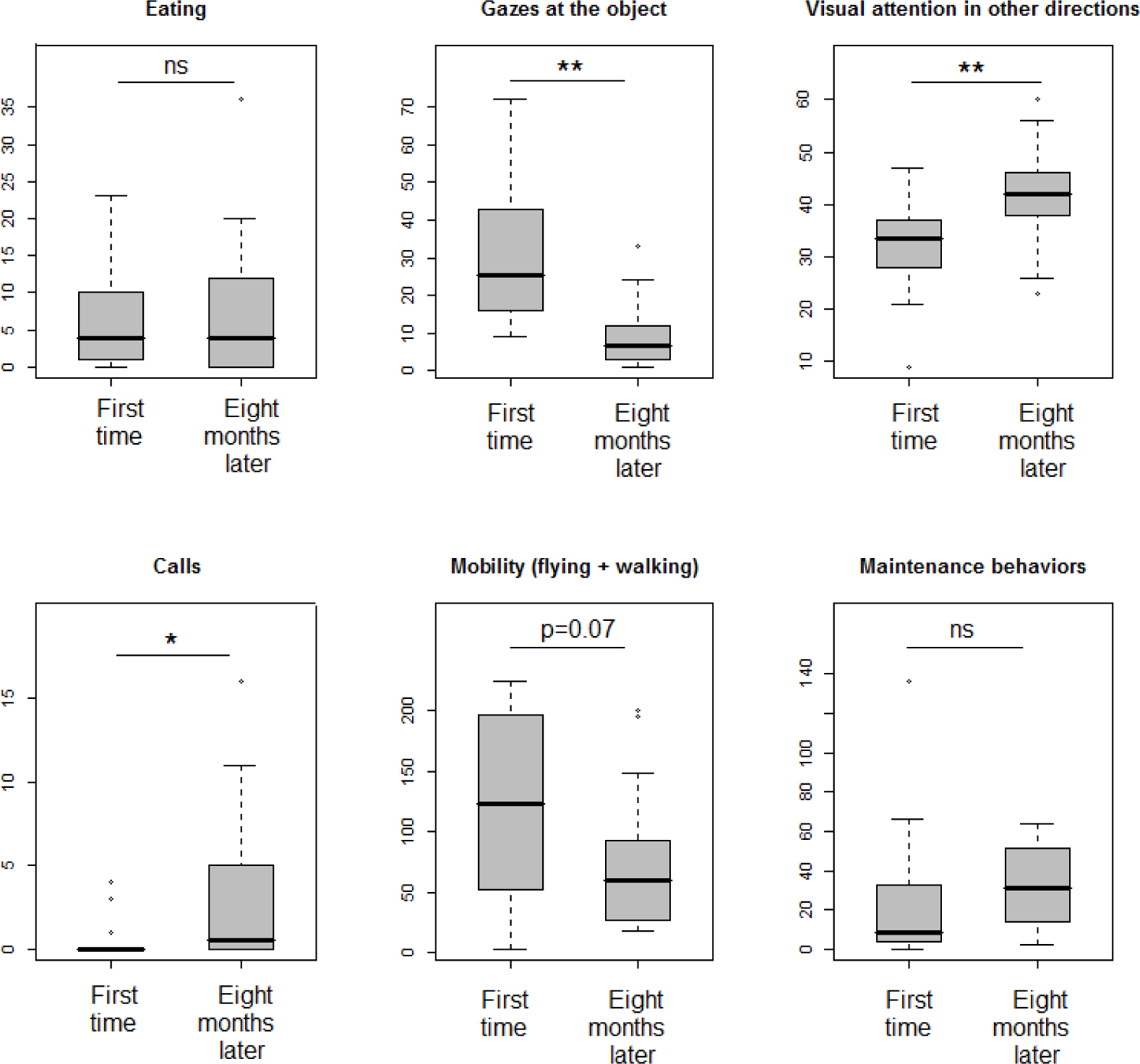
Time and attempts performance in the learning task

It is important to note that during the experiment to test for memorization of the object some individuals rushed on the object and looked inside very probably searching for a worm even if their was no worm in the object memorization test. That probably means that eight month later there were individuals who remembered the past presence of worms in the box.

## Discussion

### Habituation in starlings

We observed different types of behavioural plasticity in the experiments we conducted. Almost all the adult and young starlings confronted with a novel object did not touch it in a first encounter of 15 minutes with it. However, after 24h of presence of the object and the addition of a flour worm in it, almost all the birds touched the object and tried to open the box.

Habituation and motivation seem to act together as enhancers to approach the new objects ; they may replace or counterbalance the preliminary neophobia. This is an example of plastic behaviour in this species and is in agreement with the theory of behavioural flexibility in invasive species (Sol et. al 2002)

Starlings appeared to be able to reduce their flight reactions when confronted successively to two different novel objects. That means that habituation is not restricted to the successive encounters of a same object but that a second new object of different color and shape may induce a wicker fear reaction when the individual has already been confronted to another novel object before. Other authors have also observed these kinds of phenomena. For example, Fox and Millam (2007) have found in Amazon parrots, *Amazona amazonica*, that individuals who had experienced various novel objects in the past expressed less neophobic reactions when confronted to other novel objects. Jones (1986) has also observed this phenomenon concerning novel food. Chicks who had been confronted with food with various colors show less neophobic reactions when confronted later to an unfamiliar blue food.

Several studies on environment enrichment in rodents had obtained the same kind of results as individuals previously reared in enriched environments containing various different objects are less neophobic than individuals reared in poor ones (Huck et al.1975, Chapillon et al. 1999, Zimmermann et al. 2001). However, the effects of enrichment on neophobia and later explorative behaviours depend also on the types of enrichment used (objects’ colors and forms), on the frequency of individuals exposure to them and on individuals’ personality (Jones 2000, Fox and Millam 2007). The use of big or fearful objects, or the too frequent use of novel objects may on the contrary accentuate neophobia.

If we translate these observations to the field of biological invasions we can suggest that depending on the early experience of individuals and on the kind and the frequencies of objects they were exposed to, individuals would vary in their probability to initiate behavioural approaches towards novel objects in the new colonized areas. Moreover, we observed that flight reactions were reduced between the first and the second object even if they were presented on two successive days. The second object was taller than the first, so even if we can think that changes in neophobia were due to changes in the shape of the objects the second one should probably elicit more fear reactions. This observation probably indicates that if an individual arrives into an environment containing various types of objects he would probably be less and less stressed by novel objects even if the second ones would a priori be more scaring. The ontogenetic situation of encountering successive novel objects may act as an enhancer of invasion process by habituation or as an accelarator during categorization of various objects in non dangerous items. This hypothesis is supported by the fact that the frequency of times gazing at the object on the two successive different objects was maintained, indicating that even if the second object elicited less flight reactions it has also provoked equal visual attention in the birds, they did not ignore it. Moreover, it has been demonstrated that starlings are able to react differently to familiar and unfamiliar songs and their neuronal and behavioural responses argue for categorization processes (Hausberger and Cousillas 1996, Hausberger and al. 1997, Hausberger and al. 2000). During the experiments we conducted, we were probably watching a categorization process in which starlings distinguished two different objects but were probably able to generalized the two situations and categorize each object in a different category by they form but in a same category of non dangerous objects. The fact that visual attention was similar between object 1 and object 2 but that visual attention significantly decreased in the memorization experiment when the same object 1 was presented eight months later converges also with the probability of a categorization process as it has been demonstrated in human babies (Fantz 1964). In human babies visual attention is the same between new images but decreases when the images are not new anymore for the baby.

It is also possible that the fact that the two objects had a same white wooden support could have been used as a common element acting as a visual cue helping to decrease flights reactions in the tested birds as categorization can be done by common elements between the visual items (Rakison and Butterworth, 1998). However, more observations should be conducted in starlings with other objects and a randomization of the order of presentation of the new objects to what individuals are exposed to.

### Solving task

In our experiments we observed that almost all the individuals where able to perform the task consisting in opening the box. It has been demonstrated that this species is able to solve different kinds of problems, to learn by association and to learn by social facilitation (Lejeune 1980, Boogert et al. 2006, 2008). Here we report that the ability to solve problems is conditioned in this species by time to reduce neophobia and by the abilities in manipulating objects with the bill and legs. Boogert and collaborators (2006) found no correlation between the latency to feed near a novel object and the learning performance of individuals. They suggested that interindividual differences in learning performance were due to learning abilities. In our case the time to open the box was explained by time to touch it and by time to manipulate it in the group of rural young but it was only explained by the time to touch the object in the rural young group. For the group of migratory adults the factor explaining the time to open the box was the manipulation time. The rural group was the one who improved more in the task and adult migratory took longer to improve because of their ability and their neophobia. Almost all the birds were able to open the box and to improve the task in at least one of the successive trials. We suggest that in invasion processess species like the European starling habituate and learn quickly how to solve problems by manipulating objects. Young birds may play a key role here because of their neophilia and tendency to explore and play with novel objects (Campbell et al. 1999). The more curious individuals may develop skills sooner and act as leaders for others who can learn by joining and observing them (Fawcett et al. 2002, Rodriguez et al. 2010). In other invasive taxa as in corvids it has been demonstrated that there are explorative innovators and that birds may use social information comming from different demonstrators (Miller et al. 2016).

### Memorization

Finally, we report here for the first time the existence of a long term memory in young starlings as they were able to remember an object seen eight months before. Young starlings leave the nest when they are 20-25 days old. During the first days out, they follow their parents and learn how to capture worms in the ground. Then they form big groups of young and fly away from their original area. During the summer they are very erratic and realize the young dispersal. At this moment they may enter very different areas previously unknown for them. This dispersal acts as an exploratory phase during which the young prospect and may learn about suitable feeding or breeding sites. The following year some of them come back to their original area or go to some of the suitable prospected areas. Having a long term memory may facilitate this return to suitable places.

Concerning migratory populations, individuals leave and return to their original areas every year. In both cases (erratic sedentary young or migratory starlings) birds may need to memorize some of the habitat elements. Mettke-Hofmann and Gwinner (2003) demonstrated with hand-reared birds that long term-spatial memory (of at least one year) exists in a migrant warbler species and is absent in an non-migrant one. Our results are in accordance with their observations and support the hypothesis of the existence of a long term memory in an invasive and migratory bird like starling ; several of the subjects tested remember the object and how to open the box. It appears that particular behaviours can be mantained across the time and constitute an advantage to exploit food resources when going to settle in a habitat that could have been prospected some seasons before. Memorization and use of social peers’ song has been demonstrated in starlings. Individuals share the song of their social peers and when they are removed from their original social group and put with new conspecifics they tend to loose their ancient repertoire and to integrate new songs shared with their new social partners (Hausberger 1993, Hausberger et al. 1995). Our observations argue for plasticity in memory which can be an adaptive attribute to permanent an changing environments.

### Effects of novelty and reinforcement on learning and memorization

Both the object and the solving task were new for the starlings when they were confronted to them at the age of seven months. It has been reported that in humans and rodents observing something new facilitates learning and memorization just after seeing novel stimuli like images or when exploring a novel environment (Davis et al. 2004, Fenker et al. 2008). It seems that being confronted to something new enhances long-term potentiation and facilitates information storage by the hippocampus which increases the probability of a memory formation (Davis et al. 2004).

This mechanism may explain the memorization of the object in the starlings we studied. Individuals that leave their original habitat (invaders or migratory) are confronted to many different new situations (novel spaces, novel foods, novel objects, novel conspecifics or heterospecifics) and need to integrate the numerous new information they are confronted with. It has been demonstrated that hipocampal synaptic activity allows storage information.

In mammals, long-term potentiation in situations involving novelty occurs in a particular area of the hippocampus: the dentate gyrus (Davis et al. 2004, Kitchigina et al. 2006). This structure has not been reported in birds but there is probably an analogous structure in their hippocampus acting in these kinds of processes. However hippocampus’ role in memory formation in birds has been reported in food storage and migratory species (Krebs et al. 1989, 1996, Shapiro and Wierazko 1996). Such mechanisms may be also involved in memorization of novel situations in colonization processes.

Finally, concerning positive reinforcement effects of the worm we put in the box we may suggest that the fact that individuals were deprived of food before the learning experiment has probably facilitated learning and raised the relative importance of the reinforcement. Memories associated with rewards are probably long-term maintained easier. Marsh et al. (2004) have observed that the energetic state of an animal at the moment he is learning about a food source affects the likelihood that the source will be selected in future foraging situations. In their study, starlings had to peck on keys with different colors to obtain equal quantities of food depending on if they were deprived or not of food before the experiment. When they were confronted to the different keys in subsequent observations birds preferred pecking on the keys they had been confronted with during the experiment under food deprivation than when not deprived.

## Conclusion

Our observations are in agreement with the hipotesis of behavioural plasticity and flexibility that may explain biological invasions. Modulation of neophobia and the reduction of emotional reactions by habituation enhance the long term tolerance of novel objects in an invasive bird. Explorative behaviour facilitates task solving in contexts involving novel objects and food. This aspect is exacerbated in young individuals. Finally, cognitive processes as categorisation and memorisation allow information storage which can be usefull in migration periods or when revisiting potential suitable sites during colonisation processess.

## Notes

### Competing Interest Statement

The authors have declared no competing interest.

## References

1. Boogert, N.J., Reder, S.M., & Laland, K. (2006). The relation between social rank, neophobia and individual learning in starlings. Animal Behaviour 72, 1229–1239.

2. Boogert, N.J., Reader, S.M., Hoppit, W., & Laland, K. (2008). The origin and spread of innovations in starlings. Animal Behaviour 75, 1509–1518.

3. Burger, J., & Gochfeld, M. (1983). Behaviour of nine avian species at a Florida garbage dump. Colonial Waterbirds 6, 54–63.

4. Campbell, F.M., Heyes, C.M., & Goldsmith, A.R. (1999). Stimulus learning and response learning by observation in the European starling, in a two-object/two-action test. Animal Behaviour 58, 151–158.

5. Chapillon, P., Manneché, C., Belzung C., & Caston, J. (1999). Rearing environmental enrichment in two inbred strains of mice, 1. Effects on emotional reactivity. Behaviour genetics 29, 41–46.

6. Davis, C.D., Jones, F.L., & Derrick, B.E. (2004). Novel environments enhance the induction and maintenance of long-term potentiation in the Dentate Gyrus. The Journal of Neuroscience 24, 6497–6506.

7. Echeverria, A.I., Vasallo, A. I., & Isacch, J.P. (2006). Experimental analysis of novelty responses in a bird assemblage inhabiting in suburban marsh. Canadian journal of Zoology 84, 974–980.

8. Fenker, D., Frey, J., Schuetze, H., Heipertz, D., Heinze, H-J, & Duzel E.. (2008). Novel Scenes improve recollection and recall of words. Journal of Cognitive Neuroscience. 20(7), 1250–1265.

9. Fawcett, T.W., Skinner, A.M.J., & Goldsmith, A.R. (2002). A test of imitative learning in starlings using a two-action method with an enhanced ghost control. Animal Behaviour 64, 547–556.

10. Fisher, J., & Hinde R.A.(1949). The opening of milk bottles by birds. British Birds 42,347–357.

11. Fox, R.A, & Millam R. (2004). The effect of early environment on neophobia in orange- winged Amazon parrots (*Amazona amazonica*). Applied Animal Behaviour Science 89, 117–129.

12. Fantz, R.L. (1964). Visual experience in infants, Decreased attention familiar patterns relative to novel ones. Science, 146,668–670.

13. Gadjon, G.K., Fijn, N., Huber, & L. (2006). Limited spread of innovation in a wild parrot, the kea (*Nestor notabilis*). Animal Cognition 9, 173–181.

14. Greenberg, R. (2003). The role of neophobia and neophilia in the development of innovative behaviour of birds. In Animal Innovation. Oxford University Press. Ed. Simon Reader and Kevin Laland p. 344.

15. Heffernan, D. Andelt, W., & Shivik J.A. (2005). Coyote investigative behaviour following removal of novel stimuli. The Journal of Wildlife Management 71, 587–593.

16. Hausberger, M. (1993). How studies on vocal communication in birds contribute to a comparative approach of cognition. Etologia 3, 171–185.

17. Hausberger, M., Richard-Yris, M.A., Henry, L., Lepage, L., & Schmidt, I. (1995). Song sharing reflects the social organisation in a captive group of European starlings (*Sturnus vulgaris*). Journal of Comparative Psychology 109, 222–241.

18. Hausberger, M., Richard-Yris, M.A., Henry, L., Lepage, L., & Schmidt, I. (1995). Song sharing reflects the social organisation in a captive group of European starlings (*Sturnus vulgaris*). Journal of Comparative Psychology 109, 222–241.

19. Hausberger, M., & Cousillas H. (1996). Categorization in birdsong, from behavioural to neuronal responses. Behavioural Processes 35, 83–91.

20. Hausberger, M., Forasté, M., Richard-Yris, M.A., & Nygren, C. (1997). Differential response of female starlings to shared and nonshared song types. Ethologia 5, 31–38.

21. Hausberger, M., Leppelsack E., Richard J.P., & Leppelsack, H. (2000). Neuronal bases of categorization in starling song. Behavioural Brain Research 114, 89–95.

22. Huck, U.U., & Price, E.O.(1975). Differential effects of environmental enrichment on the open-field behaviour of wild and domestic Norway rats. Journal of Comparative and Physiological Psychology 89, 892–898.

23. Jones, B.R. (1986). Responses of domestic chicks to novel food as a function of sex, strain and previous experience. Behavioural Processes 12, 261–271.

24. Jones, R.B., Carmichael, N.L., & Rayner, E. (2000). Pecking preferences and pre- dispositions in domestic chicks, Implications for the development of environmental enrichment devices. Applied Animal Behaviour Science 69, 291–312.

25. Kitchigina, V., Vankov, A., Harley, C., & Sara S.J. (2006). Novelty-elicited, noradrenaline- dependent enhancement of excitability in the dentate gyrus. European Journal of Neuroscience 9, 41–47.

26. Krebs, J.R., Sherry, D.F., Healy, S.D., Perry, V.H., & Vaccarino, A.L. (1989). Hippocampal specialization of food-storing birds. Proceedings of the National Academy of Science 86, 1388–1392.

27. Lejeune, A. (1980). Acquisition d’une réponse discriminative chez l’Etourneau (Sturnus vulgaris) en volière. Le Gerfaut 70 , 481–485.

28. Marsh, B., Schuck-Paim, C., & Kacelnik, A. (2004). Energetic state during learning affects foraging choices in starlings. Behavioural Ecology 15, 396–399.

29. Mettke-Hofmann, C., Winkler, H., & Leisler, B. (2002). The significance of ecological factors for exploration and neophobia in parrots. Ethology 108 (3), 249–272.

30. Mettke-Hofmann, C., & Gwinner, E. (2003). Long-term memory for a life on the move. PNAS 100, 5863–5866.

31. Miller, R., Schwab, C., & Bugnyar, T. (2016). Explorative Innovators and Flexible Use of Social Information in Common Ravens (*Corvus corax*) and Carrion Crows (*Corvus corone*). Journal of Comparative Psychology 130(4):328–340.

32. Pell, A.S., & Tidemann C.R. (1997). The impact of two exotic hollow-nesting birds on two native parrots in savannah and woodland in eastern Australia. Biological Conservation 79: 145–153.

33. Peris, S., Soave, G., Camperi, A., Darrieu, C., Aramburu, R. (2005). Range expansion of the European starling *Sturnus vulgaris* in Argentina. Expansion del Estornino pinto Sturnus vulvaris en Argentina. Ardeola 52: 359–364.

34. Rakinson, D.H., & Butterworth, G.E. (1998). Infants’ use of object parts in early categorization. Developmental Psychobiology 34, 49–62.

35. Rodríguez A, Clergeau P. Hausberger M. 2010. Flexibility in the use of social information in the successful *Sturnus vulgaris*: experiments with decoys in different populations. Animal Behavior 80:965–973.

36. Rodriguez, A, Gasc, A, Pavoine, S, Grandcolas, P, Gaucher, P, & Sueur, J. (2014). Temporal and spatial dynamics of animal sound within a neotropical forest. Ecological Informatics 21, 133–143.

37. Rodriguez, A., Hauberger, M., Henry, L., Clergeau, P. (2018). Differences in neophobia in an invasive bird depending on its colonization history. Submitted

38. Seferta, A., Guay, P-J., Marzinotto, E., & Lefebvre, L. (2001). Learning differences between Feral Pigeons and Zenaida Doves, the role of neophobia and human proximity. Ethology 107, 281–293.

39. Shapiro, E., & Wierazko A. (1996). Comparative, in vitro, studies of hippocampal tissue from homing and non-homing pigeon. Brain Research 725, 199–206.

40. Sol, D., Timmermans, S., Lefebvre, L. (2002). Behavioural flexibility and invasion success in birds. Animal Behaviour 63: 495–502.

41. Zimmermann, A., Stauffacher, M., Langhans, W., Würbel, H. (2001). Enrichment-dependent differences in novelty exploration in rats can be explained by habituation. Behavioural Brain Research 121, 11–20.

